## Supplemental Figures for "Seed structure and phosphorylation in the fuzzy coat impact tau seeding competency"

**Table S1**

| Antibody | Vendor | Host | Catalog Number | Dilution | Use |
|---|---|---|---|---|---|
| Anti-Streptomyces griseus Pronase | LS Bio | Rabbit | LS-C147534 | 1:2,000 | WB |
| Anti-Tau, Clone T49 | Millipore Sigma | Mouse | MABN827 | 1:1,500 | ICC |
| AT8 | Thermo Fisher | Mouse | | 1:1,000<br>1:500 | IHC<br>WB |
| Goat anti-Mouse IgG (H+L) Highly Cross-Adsorbed Secondary Antibody, Alexa Fluor™ Plus 680 | Thermo Fisher | Goat | A32729 | 1:10,000 | WB |
| Goat anti-Rabbit IgG (H+L) Highly Cross-Adsorbed Secondary Antibody, Alexa Fluor™ Plus 680 | Thermo Fisher | Goat | A32734 | 1:10,000 | WB |
| Goat anti-Rabbit IgG (H+L) Highly Cross-Adsorbed Secondary Antibody, Alexa Fluor™ Plus 800 | Thermo Fisher | Goat | A32735 | 1:10,000 | WB |
| Goat anti-Mouse IgG (H+L) Highly Cross-Adsorbed Secondary Antibody, Alexa Fluor™ Plus 800 | Thermo Fisher | Goat | A32730 | 1:10,000 | WB |
| Alexa Fluor 546 Goat Anti-Mouse IgG (H+L) | Thermo Fisher | Goat | A11003 | 1:500 | ICC |
| Goat Anti-Rabbit IgG Biotinylated | Vector Laboratories | Goat | BA-1000-1.5 | 1:1,000 | IHC |
| PHF1 (Supernatant) | Peter Davies | Mouse | AB_2315150 | 1:250 | WB |
| Phospho-Tau Ser262 | Thermo Fisher | Rabbit | 44-750G | 1:500 | WB |
| Recombinant Anti-Mouse IgG1 | Abcam | Rabbit | ab190481 | 1:1,000 | IHC |
| Recombinant Anti-Tau EP2456Y | Abcam | Rabbit | ab76128 | 1:2,000 | WB |
| Tau 640-680 | Osenses | Rabbit | OST00329W | 1:2,000 | WB |
| Tau5 | Binder/Kanaan | Mouse | AB_2721194 | 1:500 | WB |

**Table 1.** Table of antibodies used. Abbreviations: Immunohistochemistry (IHC), Western Blot (WB), Immunocytochemistry (ICC)

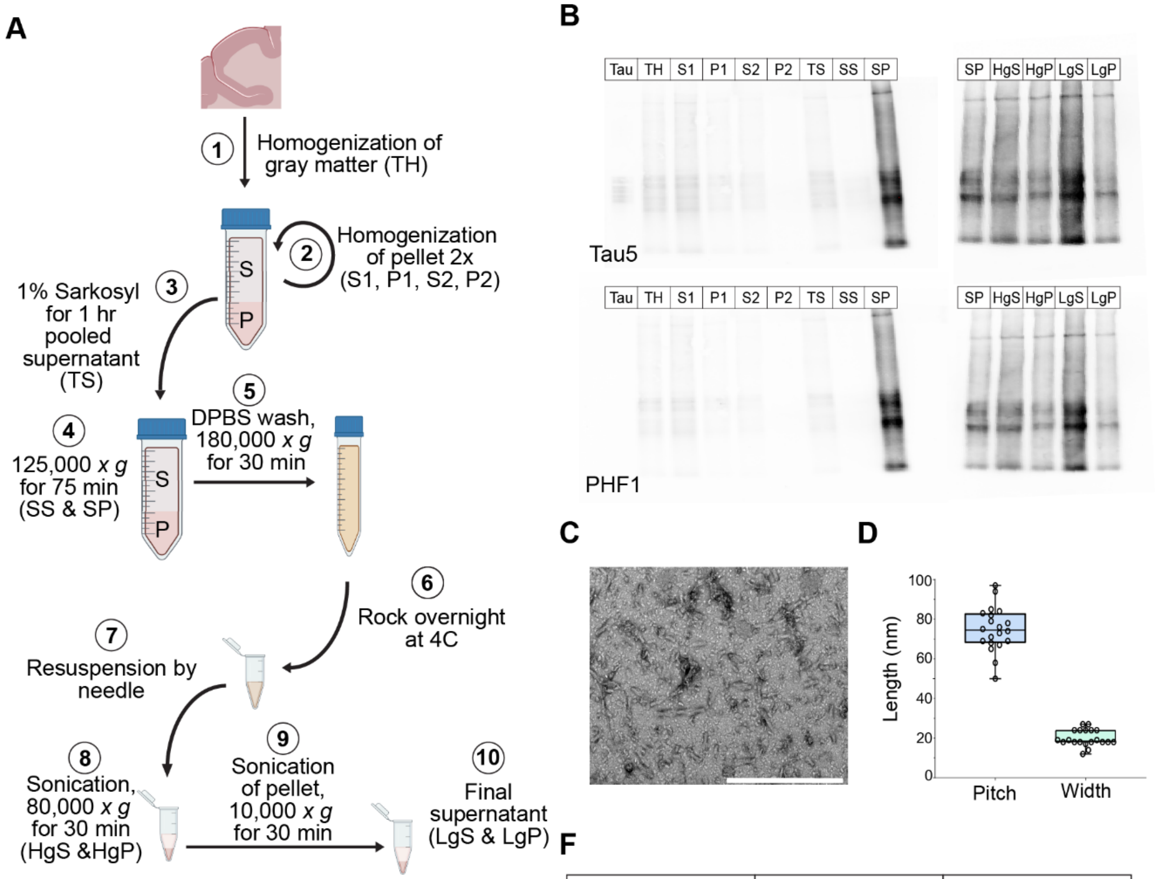


**E**

| Total Protein | Tau Protein | Tau Percentage |
|---|---|---|
| 13.4 mg/mL | 2.8 mg/mL | 20.9% |

**F**

| Age | Sex | PMI |
|---|---|---|
| 77 | F | 15.1 hours |
| 65 | F | 22 hours |
| 75 | F | 17.14 hours |

**Figure S1. Preparation and characterization of AD Tau.** (A) Schematic of isolation of AD tau from cortical brain tissue. (B) Western blots of saved fractions from tau isolation for Tau5 (top) and PHF1 (bottom) compared to a standard tau ladder (Tau). TH – total homogenate, S1 – supernatant 1, P1- pellet 1, S2 – supernatant 2, P2 – pellet 2, TS – total supernatant, SS – sarkosyl supernatant, SP – sarkosyl pellet, HgS – high *g* spin supernatant, HgP – high *g* spin pellet, LgS – low *g* spin supernatant, LgP – low *g* spin pellet. (C) Representative TEM image of isolated AD tau used in this study. Scale bar 0.5 µm. (D) Measured width and pitch of AD tau fibrils from 20 fibrils across five images. (E) Table of protein concentration of the AD tau preparation. (F) Table of sample information for pooled brain samples used in the AD tau preparation.

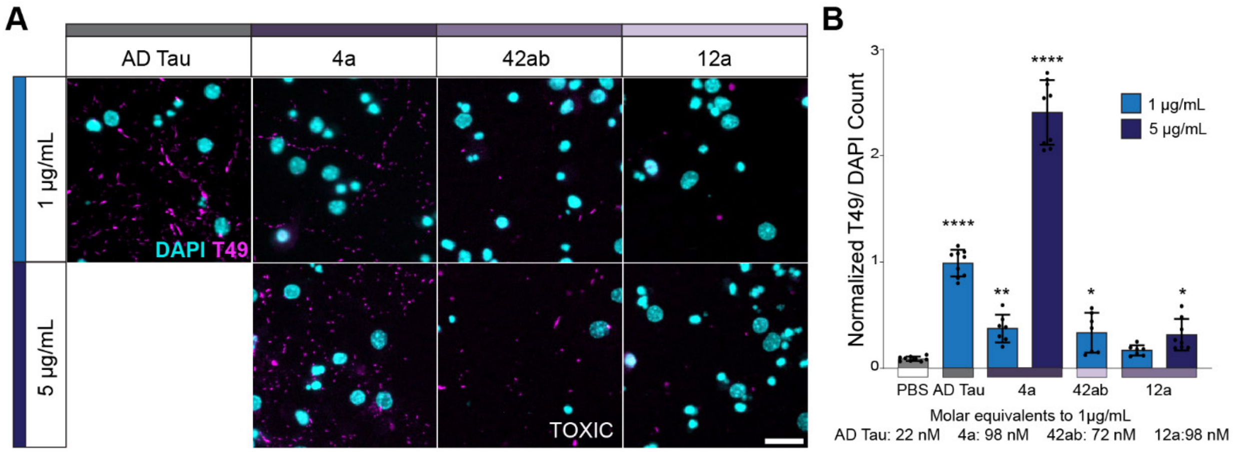


**Figure S2. Structurally-defined recombinant tau fibrils induce differential pathology in primary hippocampal neurons.** (A) Representative images of primary hippocampal neurons treated with sonicated tau fibrils at 2 doses and maintained in culture for 21 days after fibril treatment. Scale = 20 μm. (B) Quantification of tau pathology, measured by T49, relative to DAPI count. Relative molar concentrations shown below graph. Data is presented as mean ± SEM with individual values plotted. N= 9 independent wells from 3 separate cultures. * $p < 0.05$, **** $p < 0.0001$, Welch ANOVA test and Dunnett's T3 multiple comparison test compared to PBS control.

### Figure S3

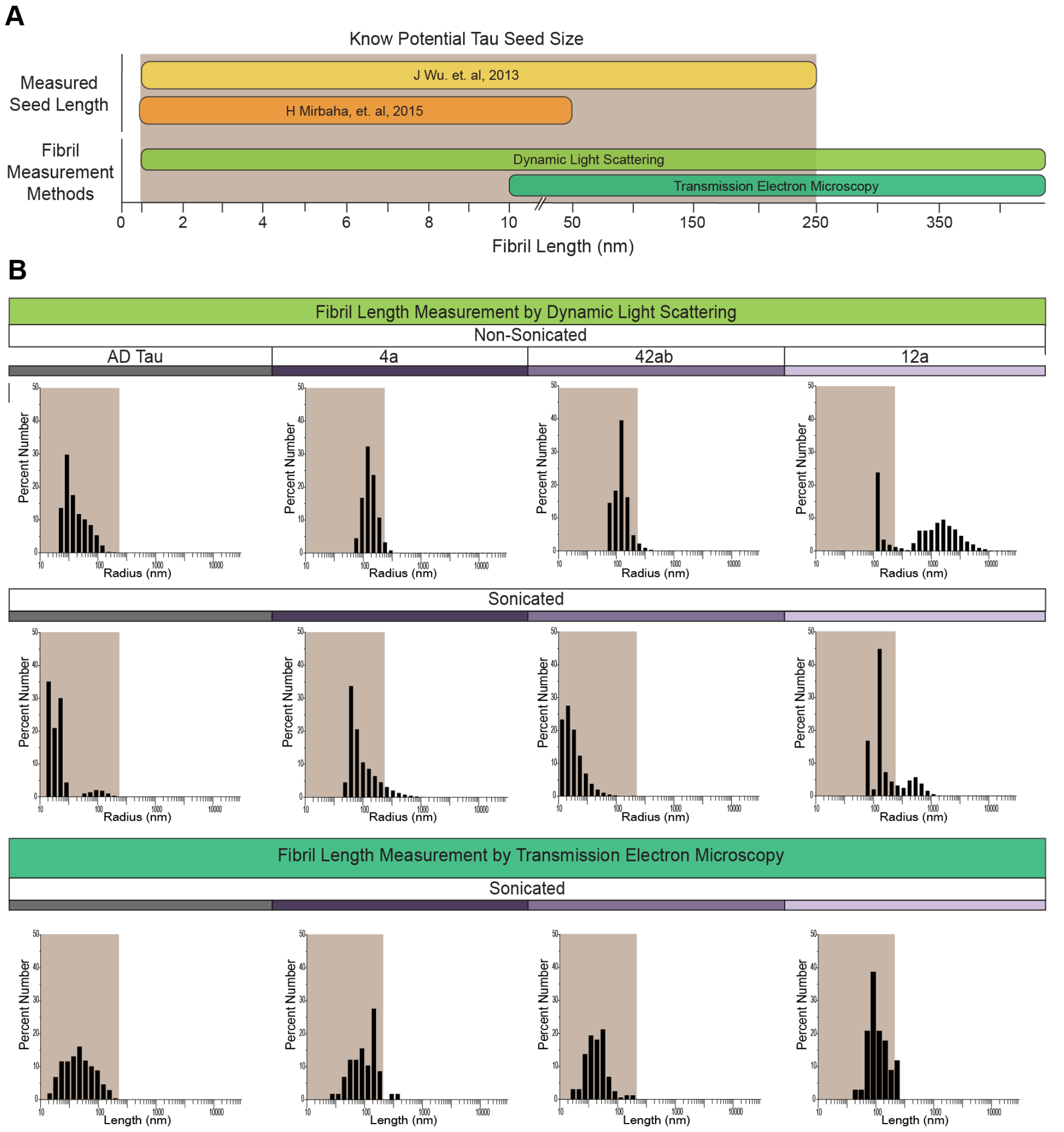


**Figure S3. Measurements of seed size in select fibrils.** (A) Graph showing known effective tau seed size from published literature and the detection range of fibril size of dynamic light scattering (DLS) and transmission electron microscopy (TEM). (B) Plotted measured fibril size from DLS for non-sonicated and sonicated fibrils and from TEM for sonicated tau fibrils. Tan box highlights the known range of effective seed size. DLS data: n=3. TEM data: PHF n = 401, 4a, n=63, 42ab = 160, 12a n= 43. Fibrils measured across 30 images from each grid with a minimum size measurement of 15 nm.

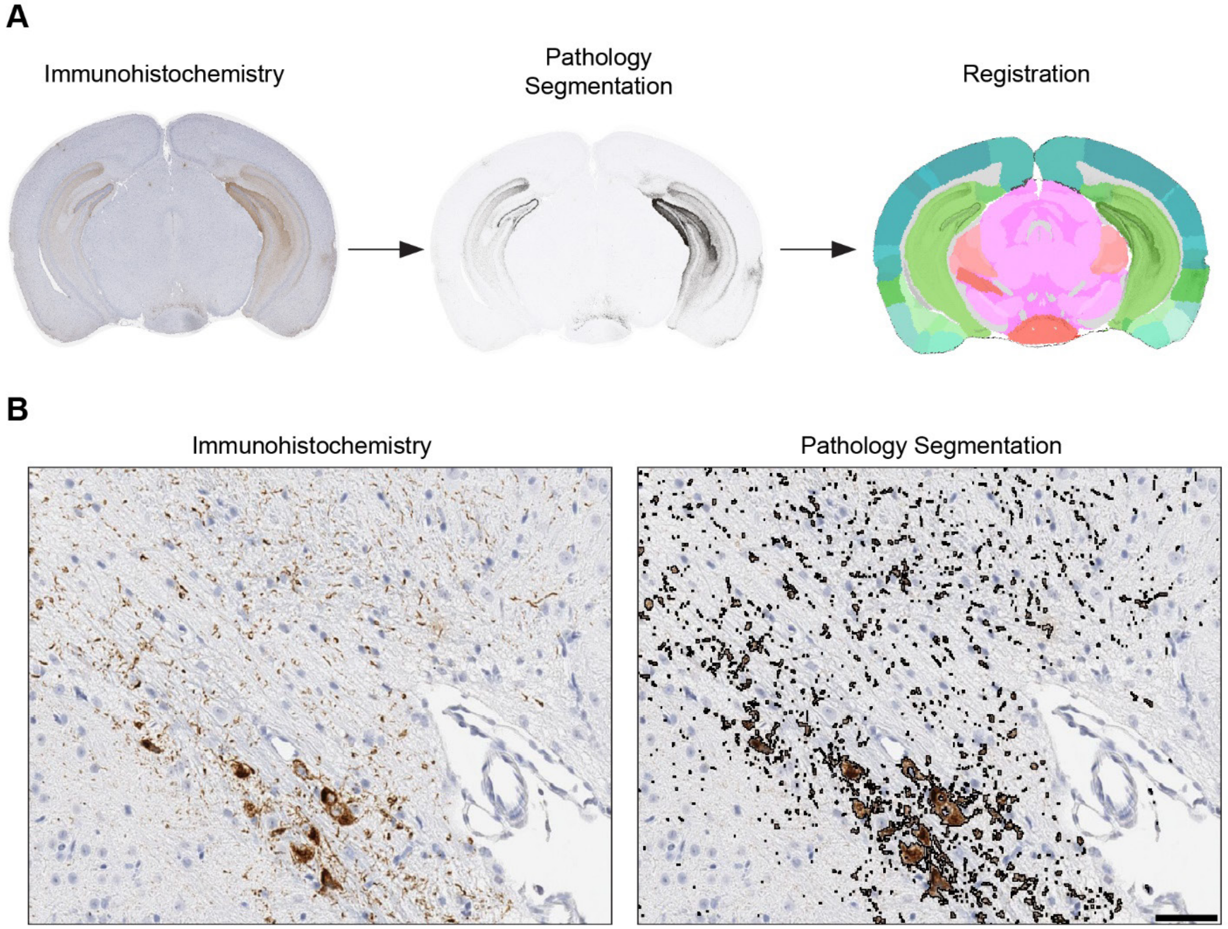


**Figure S4. Representative schematic of QUINT workflow.** (A) Representative image of immunohistochemistry stain for AT8, labeled as immunohistochemistry, pixels positive for AT8 shown in black, labeled as pathology segmentation, and an overlay of the Allen Brain Atlas to the pathology segmentation, labeled as registration. (B) Representative positive pixel detection for AT8 in the hippocampus. Scale bar = 50 mm.

**Figure S5**

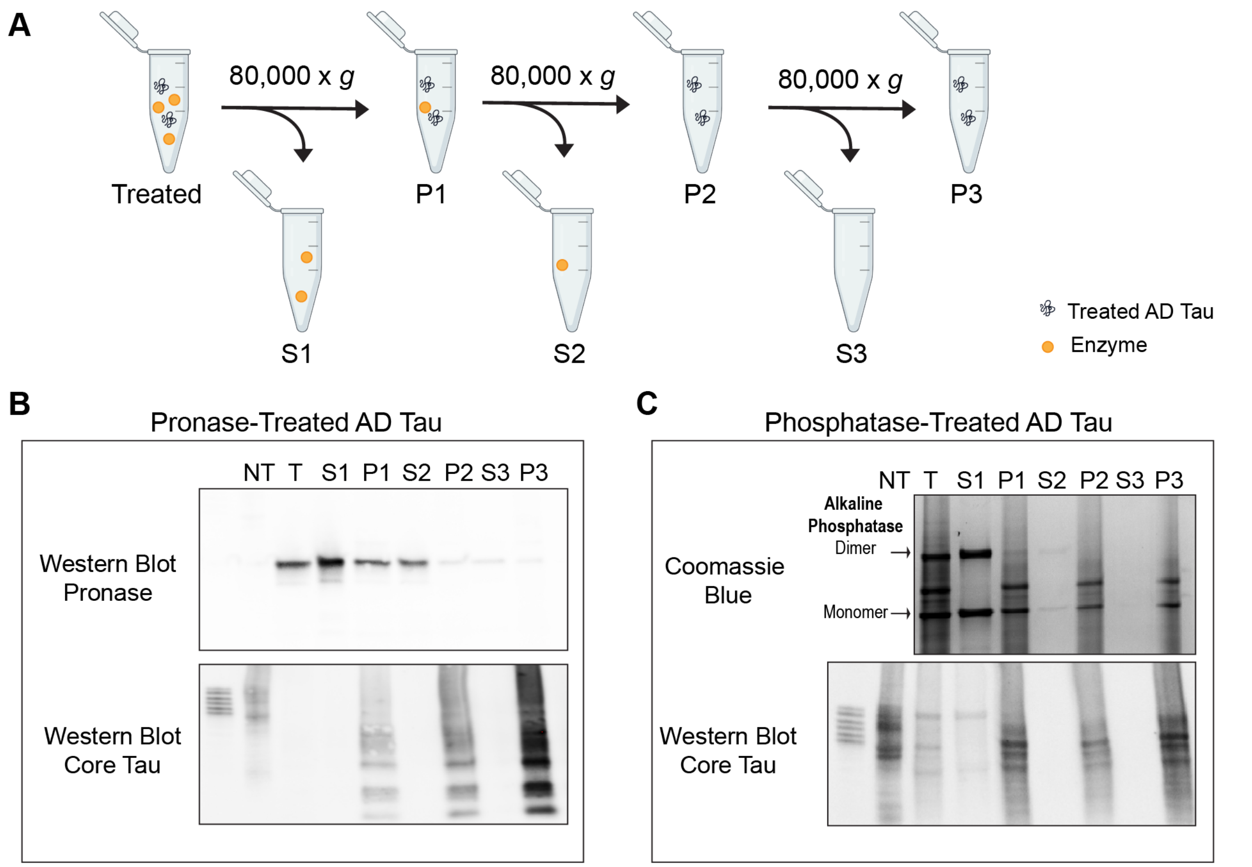


**Figure S5. Clean-up of modified tau fibrils.** (A) Schematic of removal of enzymes used in the modification of AD tau. (B) Western blot of fractions from washes of pronase-treated AD tau for pronase (top) and OST00329W (bottom). (C) Coomassie blue gel (top) and Western blot for OST00329W (bottom) of fraction from phosphatase-treated AD tau.

**Figure S6**

**Figure S6**

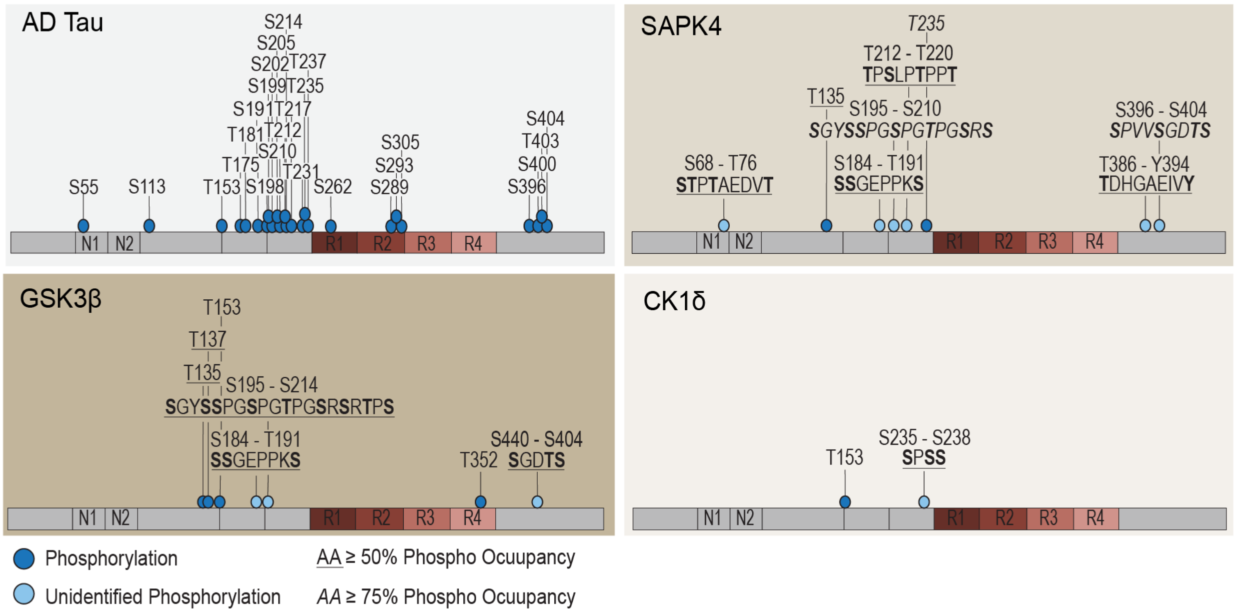


**Figure S6. Phospho-proteomic analysis following phosphorylation of tau monomer.** (A) Schematic of phosphorylation sites reported on AD PHFs[9], and phosphorylation of tau monomer by SAPK4 (top right), GSK3β (bottom left), and CK1δ (bottom right). Single phosphorylation sites are marked by a dark blue dot and an unidentified phosphorylation site is indicated by a light blue. For each unidentified phosphorylation site, the sequence is noted with the possible phosphorylated amino acid in bold. Amino acids with greater than 50% phosphor-occupancy are underlined. Amino acids with greater than 75% phosphor-occupancy are italicized. High-resolution LC-MS/MS was employed to unbiasedly identify phosphorylation sites.
